# A reference set of functional plasmids for *Vibrio natriegens*

**DOI:** 10.1101/2025.08.27.672616

**Authors:** Stephanie L. Brumwell, Ariela Esmurria, Alexander Crits-Christoph, Shinyoung Clair Kang, Charlie Gilbert, Henry H. Lee, Nili Ostrov

## Abstract

New microbial hosts with superior phenotypes, such as fast growth, are attractive for research and biotechnology, but often lack systematic evaluation of functional genetic parts. A reference set of working plasmids would increase reproducibility and encourage use of these hosts. Here, we use the POSSUM toolkit, a collection of 23 origins-of-replication and 6 antibiotic markers, to identify functional genetic parts for strains of *Vibrio natriegens*. We applied this to the wild-type strain ATCC 14048 and an engineered variant NBx CyClone™, evaluating 414 combinations of origins of replication and antibiotic selection conditions. We show that both strains support five replicons (pNG2, pSa, pSC101ts, p15A, and RSF1010) with NBx CyClone™ supporting an extra replicon RK2. The assay can be performed in under a week and is compatible with multiple DNA delivery methods. This work demonstrates the feasibility of rapidly establishing reference information to accelerate the adoption of new microbial hosts.

## Introduction

The adoption of novel microbial hosts for research and technology greatly depends on the availability and robustness of genetic tools. *Vibrio natriegens* is a promising bacterial host for accelerated cloning workflows due to its exceptionally fast growth rate^1^. It has been proposed as a superior chassis for production of proteins^2^, small molecules and materials^3–9^, as well as for DNA manipulation^10,11^.

While multiple genetic tools have been demonstrated in *V. natriegens*, information about its compatibility with existing DNA parts is a key determinant for users transitioning from other cloning methods as it greatly simplifies experimental planning and execution and increases the chances of cross-compatibility with other hosts^11–13^.

Here, we use the POSSUM toolkit^14^ to rapidly map the functionality of genetic parts, including origins of replication (ORIs) and antibiotic markers, in a wild-type strain ATCC 14048 and a commercially available strain NBx CyClone^TM^ designed for cloning.

## Results and Discussion

### ORI-marker screen

We delivered six POSSUM libraries into *V. natriegens* strains ATCC 14048 and NBx CyClone^TM^ using conjugation and chemical transformation, respectively. Each library contained a pool of 23 different ORIs and a different antibiotic selection marker: kanamycin (KAN), tetracycline (TET), chloramphenicol (CAM), gentamicin (GEN), spectinomycin (SPEC), and erythromycin (ERM) (**Supplementary Table 1, Supplementary Figure 1**). Each of the antibiotics were tested at three concentrations, for a total of 414 combinations of parts and conditions in each library experiment. The assay took three days from transformation to sequencing-ready and three days for sequencing and analysis.

We observed successful transformation in both strains using the POSSUM liquid selection assay. This assay assumes no prior knowledge regarding the optimal antibiotic concentration, and thus employs a broad range of concentrations for each antibiotic (0.25X, 1X and 4X the standard *E. coli* values)^14^ (**Supplementary Table 2**). Specifically, we observed positive hits for both strains in the liquid assay using KAN, CAM and TET libraries (**Figure 1**). The ATCC 14048 strain also showed positive hits using SPEC and ERM libraries. We could not determine GEN results for either strain, or SPEC and ERM results for NBx CyClone^TM^, due to growth of untransformed cells under selection, likely requiring further optimization of antibiotic concentrations for these strains.

Not surprisingly, our results demonstrate that altering selective conditions can affect assay sensitivity and outcome. Specifically, when using agar rather than liquid selection for the same NBx CyClone^TM^ experiment, we observed positive hits not only for KAN, CAM and TET but also for the GEN library (**Figure 2A, Supplementary Figure 2, Supplementary Figure 3**). Agar selection enabled the detection of 3 additional putative functional ORIs (RK2, pSC101ts, and pTHT15) that were not identified through liquid selection. Similarly, we also identified positive hits for GEN when using an alternative conjugation media for strain ATCC 14048 (TSB instead of PBS) (**Figure 2B**). Together, these results indicate that KAN, TET, CAM, GEN, SPEC and ERM can be used for POSSUM library selection in *V. natriegens*, and that varying selection and conjugation media can be used to increase assay sensitivity.

**Figure 1.**
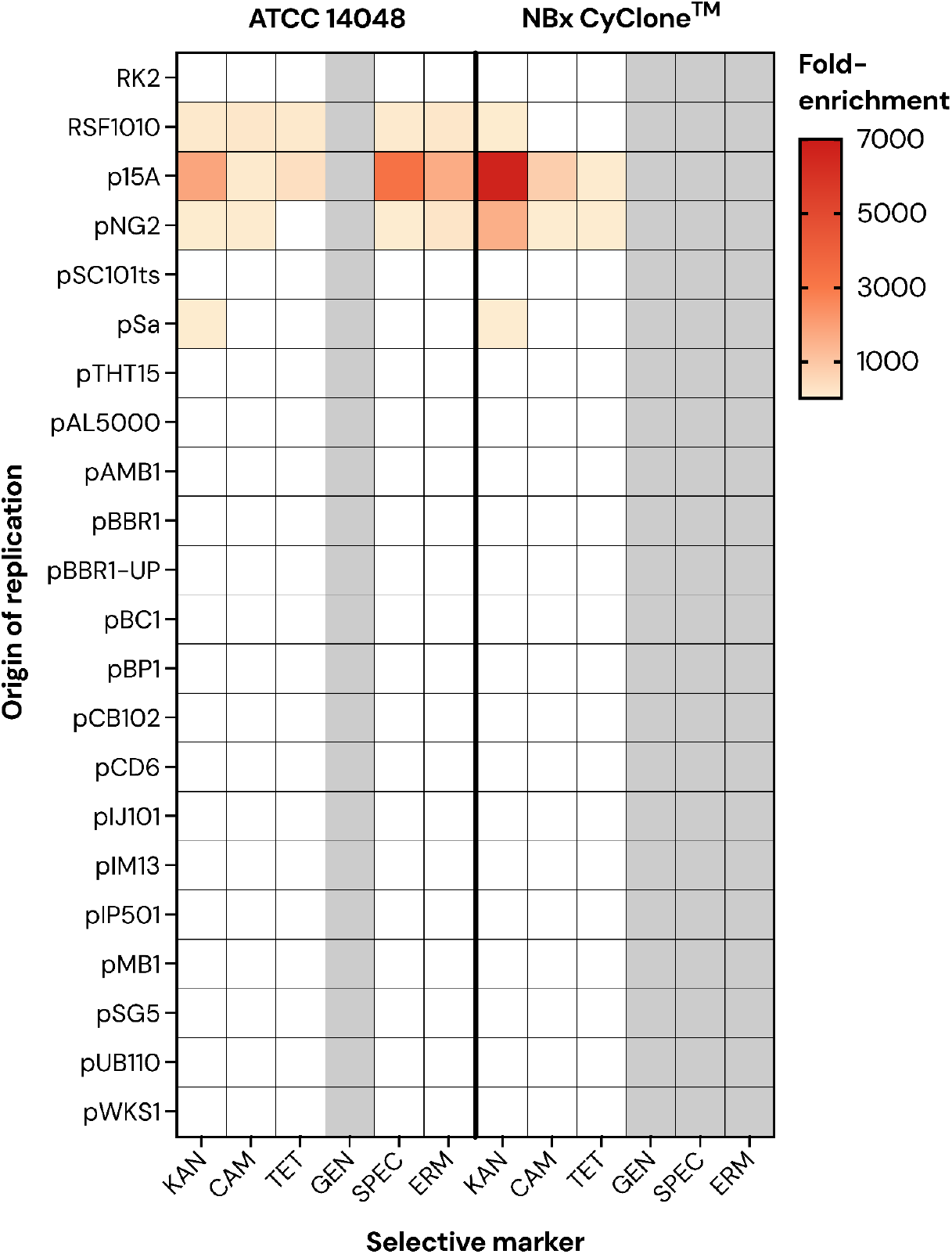
Functional origins of replication in *V. natriegens* identified by liquid selection assay. Heat map of ORIs significantly enriched over non-functional control as identified by sequencing. The same 23-ORI barcoded library was delivered into wild type (ATCC 14048) and engineered (NBx CyClone^TM^) strains using six different antibiotic selective markers. Transformants were selected in liquid media and sequenced to determine ORI enrichment. Gray boxes indicate antibiotic concentration was insufficient for selection. White boxes indicate that the ORI was not identified through sequencing.

**Figure 2.**
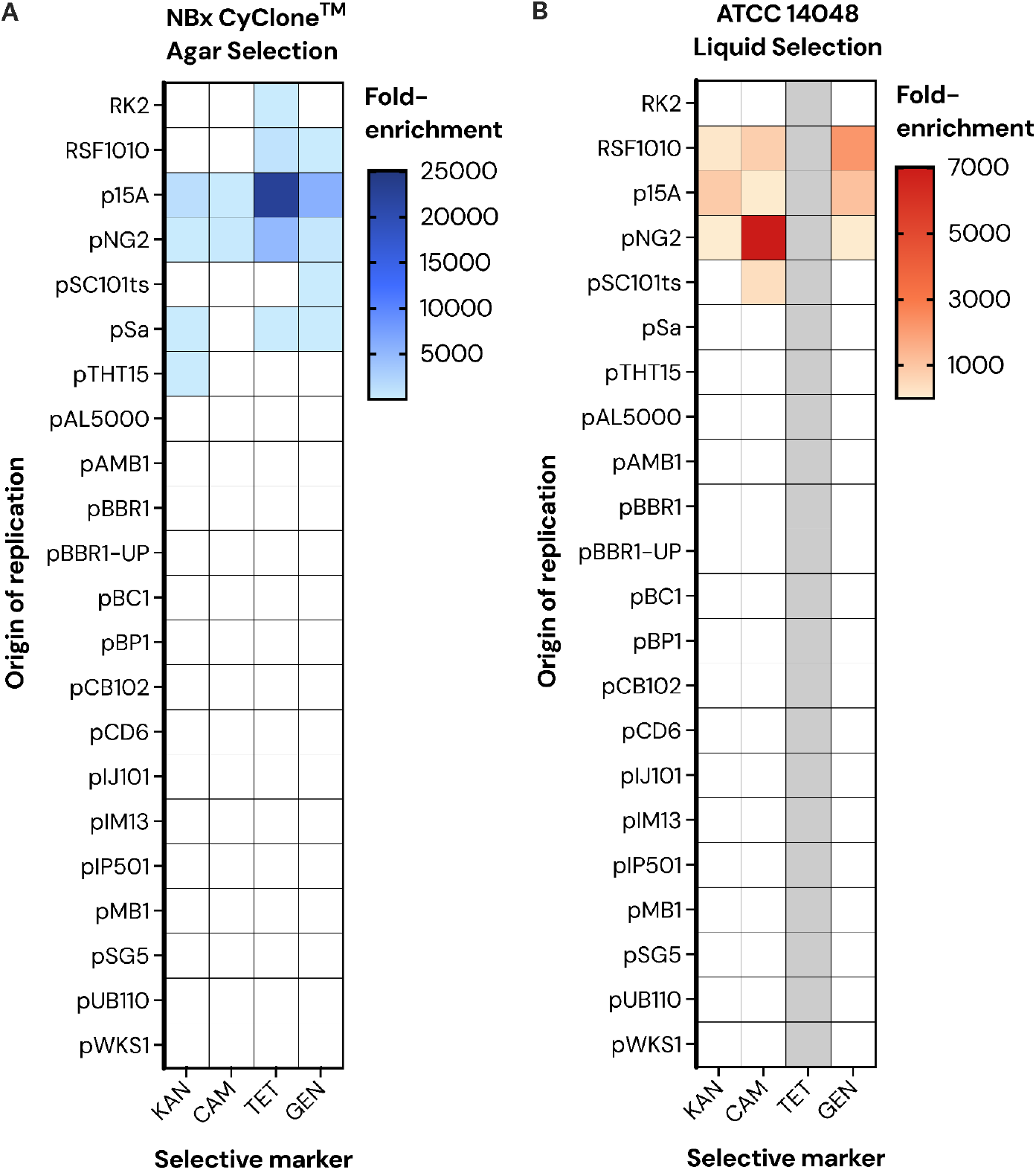
Origins of replication and selective markers identified for *V. natriegens* by altering selective or conjugative media. Heat map of ORIs significantly enriched over the non-functional control ORI following transformation of POSSUM plasmid libraries in **(A)** *V. natriegens* NBx CyClone^TM^ using solid agar selection or **(B)** *V. natriegens* ATCC 14048 using TSB conjugation media and liquid selection (see Methods). Gray boxes indicate antibiotic concentration was insufficient for selection. White boxes indicate that the ORI was not identified through sequencing.

Across all experiments, our screen identified seven putative ORIs, including three that were previously reported and noted in the NBx CyClone™ product manual (p15A, RSF1010, RK2)^12,15,16^ and four novel ORIs (pNG2, pSa, pSC101ts, and pTHT15) (**Figure 1, Figure 2, Table 1**). Notably, multiple ORIs were independently identified using different libraries, indicating replicability of POSSUM library assays. We validated the functionality of six ORIs by individually transforming each plasmid into *V. natriegens* NBx CyClone^TM^ (**Supplementary Figure 4, Supplementary Table 3**) and performed whole plasmid sequencing on novel ORIs (pNG2, pSa, pSC101ts). The pSC101 ORI has been reported previously in *V. natriegens* ATCC 14048^12^, but this is the first report on the functionality of a variant that is temperature sensitive in *E. coli*. The pTHT15 ORI, which was a low confidence hit with marginal enrichment over control (**Figure 2A**) failed the follow-up transformation and was therefore deemed nonfunctional in the NBx CyClone™ strain.

**Table 1.** *V. natriegens* origins of replication.

| ORI | This study |  | Product Manual<br>NBx<br>CyClone™ | Previously<br>Reported |
| --- | --- | --- | --- | --- |
|  | NBx<br>CyClone™ | ATCC<br>14048 |  |  |
| pMB1* | n/a | n/a | Y | 12,16 |
| pUC19** | Y | n/a | Y | 12,16 |
| p15A | Y | Y | Y | 12,16 |
| RSF1010 | Y | Y | Y | 1 |
| RK2 | Y | N | Y | 12,16 |
| pBBR1 | N | N | N | 12 |
| pNG2 | Y | Y | N | N |
| pSa | Y | Y | N | N |
| pSC101ts | Y | Y | N | N |
| pTHT15 | N | N | N | N |
| pMB1_Bifidobacterium* | N | N | N | N |
| pBBR1-UP | N | N | N | N |
| pWKS1 | N | N | N | N |
| pBP1 | N | N | N | N |
| pAL5000 | N | N | N | N |
| pAMβ1 | N | N | N | N |
| pBC1 | N | N | N | N |
| pIJ101 | N | N | N | N |
| pIM13 | N | N | N | N |
| pSG5 | N | N | N | N |
| pUB110 | N | N | N | N |
| pIP501 | N | N | N | N |
| pCB102 | N | N | N | N |
| pCD6 | N | N | N | N |

\*see Results section for pMB1 details.
\*\*pUC19 is a high copy derivative of pMB1 not included in the POSSUM toolkit. It was provided as control for the NBx CyClone™ kit. It was not tested in ATCC 14048.

Interestingly, pMB1 was described as functional in the NBx CyClone^TM^ product manual but was not identified as a hit in our screen. Comparing the sequence of pMB1 in the POSSUM kit to that used in the NBx CyClone^TM^ manual, we found that ‘pMB1’ in the scientific literature refers to different ORIs. The one described in the manual originates from *E. coli* and was the basis for pBR322/pUC19 ORIs, which are not included in the POSSUM kit^17^. Alternatively, POSSUM pMB1 was isolated from the Actinomycetota bacterium *Bifidobacterium longum*^*18*^, and it is therefore unlikely to be functional in *V. natriegens*.

The ORIs identified for NBx CyClone^TM^ align well with those we identified for *V. natriegens* ATCC 14048 using the same POSSUM libraries (**Table 1, Figure 1, Figure 2**). For *V. natriegens* ATCC 14048, RK2 and pBBR1 were not identified as functional ORIs although previous reports suggest that they should be functional^12^. The RK2 and pBBR1 ORIs were lower in abundance than the average ORI in the pool and RK2 had a lower transformation efficiency compared to the other functional ORIs identified for NBx CyClone^TM^ (**Supplementary Table 3**). These combined factors may have prevented the identification of RK2 and pBBR1 in our pooled screen.

The library was delivered by chemical transformation into NBx CyClone^TM^ and by conjugation into ATCC 14048, demonstrating the robustness of the POSSUM assay across strains and DNA delivery methods. As previously shown for other *V. natriegens* strains^1,10,12^, the transformation efficiency (TE) for NBx CyClone™ using chemical transformation was generally high with ∼10^6^ CFU/µg DNA for all libraries, between 10^3^-10^7^ for individual plasmids, and up to ∼10^8^ CFU/µg DNA using pUC19 control plasmid (**Supplementary Table 3**). Notably, we observed faster colony growth using NBx proprietary media compared with LB3, with colonies appearing on NBx media after only 6 hours (**Supplementary Figure 4**).

### Fluorescent protein expression

Since all ORI-marker plasmids possess an identical J23100-mScarlet fluorescent protein expression cassette, analysis of mScarlet fluorescence provided insights into the relative expression levels associated with the *V. natriegens* NBx CyClone^TM^ strain harboring plasmids with different ORIs. The pNG2 plasmid showed the highest mScarlet expression, followed by p15A, with minimal to no detectable expression from other plasmids (**Supplementary Figure 5**). Interestingly, for pNG2 and p15A, mScarlet expression was higher in NBx media compared to LB3 media after 24 hours of incubation, though expression in LB3 media reached similar levels after 42 hours. This suggests that pNG2 and p15A could be valuable ORIs for applications requiring high gene expression, while other ORIs might be better suited for lower copy number, stable maintenance, or the expression of toxic products.

## Conclusion

The POSSUM toolkit enables experimentalists to rapidly identify and compare functional genetic parts across species and strains. In this study, we provide a reference set of functional and non-functional ORIs and antibiotic selective markers for both *V. natriegens* ATCC 14048 and NBx CyClone^TM^. Our work overcomes common difficulties when working with new hosts, where each strain background may display variable antibiotic selection requirements or compatibility of genetic parts.

## Materials and Methods

### Strains and Growth Conditions

*Vibrio natriegens* NBx CyClone^TM^ was grown in NBx production media or LB3 media (Boston Bioproducts CAT#C-10106C), at 37°C or 30°C where specified. *Vibrio natriegens* 111 (ATCC 14048) was grown in MB media at 37°C. *Escherichia coli* pir-116 was grown in LB media at 37°C. *Escherichia coli* BW29427 was grown in LB media supplemented with diaminopimelic acid (60 µg/mL) at 37°C. Antibiotic supplementation for all strains is noted where relevant.

### Plasmid and Plasmid Library Preparation

Plasmids and plasmid libraries used in this study are summarized in **Supplementary Table 1**. Plasmid library DNA for pGL2_217, pGL2_220 and pGL2_221 was extracted from *E. coli* BW29427 grown to OD600 0.7-1.8 using the QIAprep Spin Miniprep. Plasmid library DNA for pGL2_218, pGL2_219 and pGL2_222 was extracted through Eton Bioscience Inc. or Quintara Biosciences plasmid preparation service. Plasmid and plasmid library DNA concentrations were measured using the Qubit 1X dsDNA High Sensitivity (HS) Kit on the Qubit™ Flex Fluorometer (Thermo Fisher Scientific) and normalized to 25 ng/µL. Input library distributions were estimated by amplicon sequencing described below.

### Transformation

Transformations using the Novel Bio CyClone^TM^ chemical competent cells were performed according to the manufacturer’s instructions^15^ with modifications as follows. <u>For plasmid library transformation</u>, 100 ng DNA (4 µL) was used per reaction. pUC19 (10 ng, 1 µL) provided in the kit was used as positive control. Water (no DNA) served as a negative control. Recovered transformations were diluted 1:10 and 100 µL was plated on solid selective media and into liquid selective media. Specific antibiotic concentrations used for solid and liquid selection are detailed in **Supplementary Table 2**. Agar experiments were performed on a single antibiotic concentration informed by either the manufacturer’s guidelines (KAN, CAM)^15^ or the liquid assay concentrations (GEN, TET, ERM, SPEC). LB3 media was used for all solid media selections, and NBx KAN200 plates were also used to select for the KAN library transformation. NBx production media was used for all liquid selections. <u>For individual plasmid transformation</u>, 2 µL was used per reaction (50 ng DNA total). Transformant colonies were counted manually. For transformation mixtures that had no colonies after 20 hours, 50 µL was spread plated onto LB3 KAN200 media. Additionally, 100 µL of all transformation mixtures was inoculated into LB3 KAN300 media.

### Harvesting Transformants

Transformants in liquid media were identified based on ΔOD600 of the plasmid transformation compared to the negative control. Transformants from solid agar plates were harvested by scraping the plate with 1 mL of PBS. Samples were centrifuged at 4000 x *g* for 10 minutes at 4°C to collect transformed cells, and the supernatant was removed.

### Genomic DNA Extraction

Genomic DNA from transformant pellets was extracted using the MagMAX™ Viral/Pathogen Ultra Nucleic Acid Isolation Kit, automated on KingFisher Flex. DNA concentrations were measured using the Qubit 1X dsDNA High Sensitivity Kit on the Qubit™ Flex Fluorometer (Thermo Fisher Scientific).

### Amplicon Sequencing of Plasmid Barcodes

Amplicon sequencing library preparation was performed as previously described^19^ with the following modifications. <u>For PCR 1</u>: Separate KAPA Hifi components - KAPA Hifi Fidelity Buffer, KAPA dNTP, and KAPA HiFi Pol - were used with a total reaction volume of 20 µL. Primers used can be found in **Supplementary Table 4**. PCR 1 cycling conditions: Initial denaturation at 95°C for 3 min, 17-22 cycles of denaturation at for 20 sec, annealing at 64°C for 15 sec, extension at 72°C for 15 sec, final extension at 72°C for 1 min, then cooling. Next, quantitative PCR was performed for PCR 1 using EvaGreen dye and cycling was stopped when the PCR amplification curve was in the exponential phase for each of the samples. <u>For PCR 2</u>: The PCR 1 product was diluted 10-fold and 1 µL was used as template for PCR 2 reaction (total volume: 20 µL). Separate KAPA Hifi components - KAPA Hifi Fidelity Buffer, KAPA dNTP, and KAPA HiFi Pol - were used with a total reaction volume of 20 µL. PCR 2 cycling conditions: Initial denaturation at 95°C for 3 min, 12 cycles of denaturation at 98°C for 20 sec, annealing at 70°C for 15 sec, extension at 72°C for 15 sec, final extension at 72°C for 1 min, then cooling. Expected size of final library construct is 232 bp.

We estimated the abundance of each plasmid/origin of replication (ORI) by counting the number of reads that mapped to the unique barcode for each ORI, as described in a previous study^19^. We then identified functional ORIs by their statistically significant enrichment compared to the background abundance of each ORI, which was determined from sequencing the input libraries (q<0.05, Bonferroni-corrected). Whole plasmid sequencing was performed on single colonies of transformants harbouring novel ORIs (pNG2, pSa and pSC101ts) using Plasmidsaurus Zeroprep.

### mScarlet Fluorescence Analysis

A kinetic run was performed on the Synergy H1 Plate Reader using a single colony from all successful transformations in NBx KAN300 and LB3 KAN300 media. OD600 and mScarlet fluorescence [excitation λ = 569 nm, emission λ = 610 nm] were measured at 5-minute intervals for 45 hours. The run was performed at 30^°^C to accommodate the growth of the temperature-sensitive pSC101ts ORI, using double orbital shaking.

### *Vibrio natriegens* ATCC 14048 ORI-screen

Performed as previously described^19^ with conjugation performed on PBS or TSB media supplemented with diaminopimelic acid (60 µg/mL) and selection in liquid MB media.

## Data and Resource Availability

The POSSUM toolkit is available via Addgene (POSSUM Toolkit, Kit ID #1000000234). Sequencing analysis scripts are available at https://github.com/cultivarium/ORI-marker-screen/.

## Author Contributions

S.L.B., C.G., N.O., and H.H.L. conceived of and designed the project. S.B. and A.E. performed the ORI-marker screen experiments. S.B. performed fluorescence experiments. C.K. performed DNA sequencing workflows. A.C.C. analyzed sequencing data. S.B., N.O., and H.H.L wrote the manuscript. All authors have read and approved the manuscript.

## Acknowledgements

We thank all members of the Cultivarium team for discussions throughout this project. Cultivarium acknowledges support from Schmidt Futures as a Convergent Research Focused Research Organization (FRO).

## References

1. Lee, H. H. et al. Vibrio natriegens, a new genomic powerhouse. bioRxiv 058487 (2016) doi:10.1101/058487.

2. Mojica, N. et al. Using Vibrio natriegens for high-yield production of challenging expression targets and for protein perdeuteration. Biochemistry 63, 587–598 (2024).

3. Sun, X. et al. Engineering of fast-growing Vibrio natriegens for biosynthesis of poly(3-hydroxybutyrate-co-lactate). Bioresour. Bioprocess. 11, 86 (2024).

4. Politan, R. J. et al. Establishing Vibrio natriegens as a high-performance host for acetate-based poly-3-hydroxybutyrate production. Metab. Eng. 92, 22–38 (2025).

5. Zheng, S. et al. Engineered Vibrio natriegens with a toxin-antitoxin system for high-productivity biotransformation of l-lysine to cadaverine. J. Agric. Food Chem. 73, 6113–6123 (2025).

6. Wu, F., Wang, S., Peng, Y., Guo, Y. & Wang, Q. Metabolic engineering of fast-growing Vibrio natriegens for efficient pyruvate production. Microb. Cell Fact. 22, 172 (2023).

7. Metabolic Engineering of Vibrio Natriegens for Poly(3-Hydroxybutyrate) Production.

8. Ellis, G. A. et al. Exploiting the feedstock flexibility of the emergent synthetic biology chassis Vibrio natriegens for engineered natural product production. Mar. Drugs 17, 679 (2019).

9. Erian, A. M., Freitag, P., Gibisch, M. & Pflügl, S. High rate 2,3-butanediol production with Vibrio natriegens. Bioresour. Technol. Rep. 10, 100408 (2020).

10. Lee, H. H., Ostrov, N., Gold, M. A. & Church, G. M. Recombineering in Vibrio natriegens. Synthetic Biology (2017).

11. Stukenberg, D. et al. The Marburg Collection: A Golden Gate DNA assembly framework for synthetic biology applications in Vibrio natriegens. ACS Synth. Biol. 10, 1904–1919 (2021).

12. Tschirhart, T. et al. Synthetic biology tools for the fast-growing marine bacterium Vibrio natriegens. ACS Synth. Biol. 8, 2069–2079 (2019).

13. Faber, A. et al. Expanding genetic engineering capabilities in Vibrio natriegens with the Vnat Collection. Nucleic Acids Res. 53, (2025).

14. Gilbert, C. et al. Design and construction towards a pan-microbial toolkit. bioRxiv 2024.02.23.581749 (2024) doi:10.1101/2024.02.23.581749.

15. NBx CyClone− Product Manual. (2025).

16. Weinstock, M. T., Hesek, E. D., Wilson, C. M. & Gibson, D. G. Vibrio natriegens as a fast-growing host for molecular biology. Nat. Methods 13, 849–851 (2016).

17. Hashimoto-Gotoh, T. & Timmis, K. N. Incompatibility properties of Col E1 and pMB1 derivative plasmids: random replication of multicopy replicons. Cell 23, 229–238 (1981).

18. Rossi, M., Brigidi, P., Gonzalez Vara y Rodriguez, A. & Matteuzzi, D. Characterization of the plasmid pMB1 from Bifidobacterium longum and its use for shuttle vector construction. Res. Microbiol. 147, 133–143 (1996).

19. Gilbert, C. et al. A scalable framework for high-throughput identification of functional origins of replication in non-model bacteria. bioRxiv 2023.05.19.541510 (2023) doi:10.1101/2023.05.19.541510.

