## Supplementary Figures and Tables for "A reference set of functional plasmids for *Vibrio natriegens*"

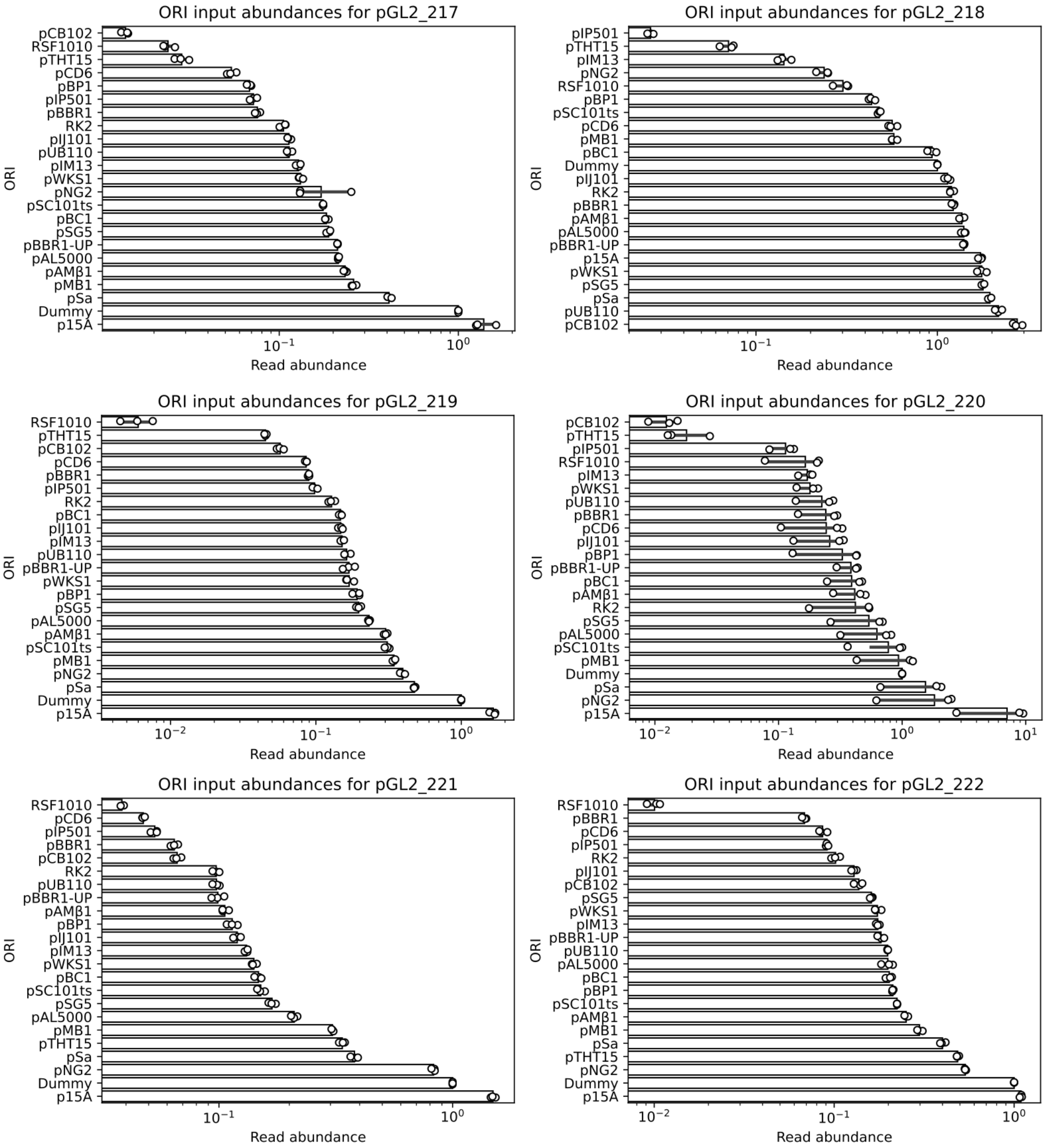


**Supplementary Figure 1. Plasmid library ORI distribution.** Distribution of the 23 origins of replication in the six POSSUM plasmid libraries: pGL2_217, KAN; pGL2_218, GEN; pGL2_219, ERM; pGL2_220, TET; pGL2_221, CAM; pGL2_222, SPEC. ORI distribution is reported as read abundances following amplicon sequencing of unique plasmid barcodes.

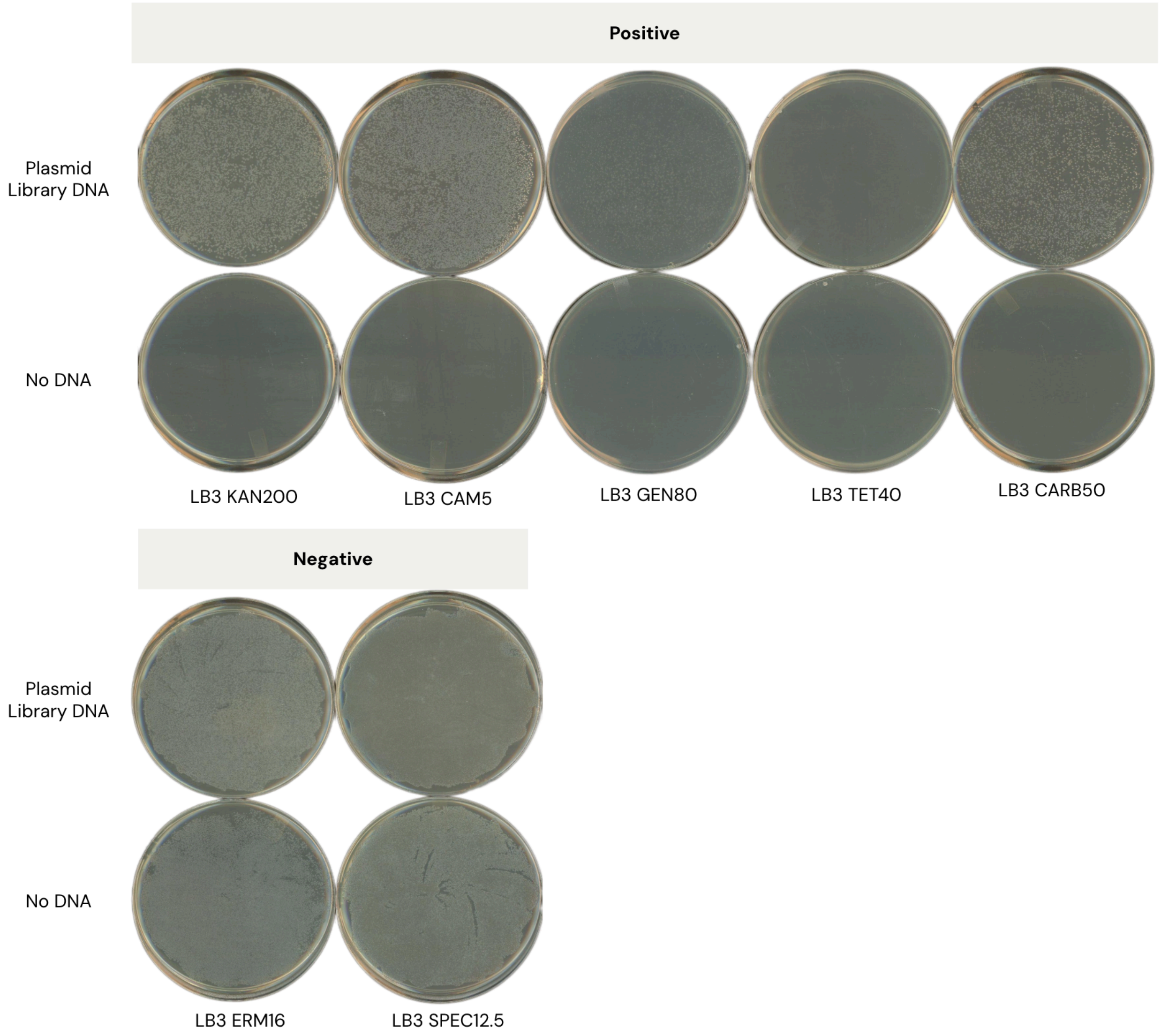


**Supplementary Figure 2. Agar selection of POSSUM libraries for *V. natriegens* NBx CyClone™.** Images of selective agar plates for *V. natriegens* NBx CyClone™ following transformation with six plasmid libraries (KAN, CAM, GEN, TET, ERM, SPEC) and a pUC19 positive control plasmid (CARB). Plates were incubated at 30°C for 20 hours. Antibiotic concentrations are reported in µg/mL (*e.g.* KAN200 refers to kanamycin 200 µg/mL).

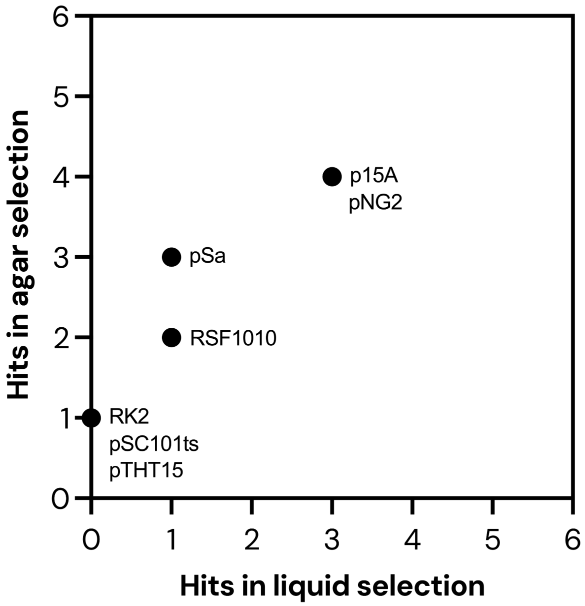


**Supplementary Figure 3. Correlation between ORI hits in liquid and agar selection for *V. natriegens* NBx CyClone™.** The number of total libraries where ORI hits were observed is shown for liquid selection (X-axis) and agar selection (Y-axis). Data points are labelled with corresponding ORIs.

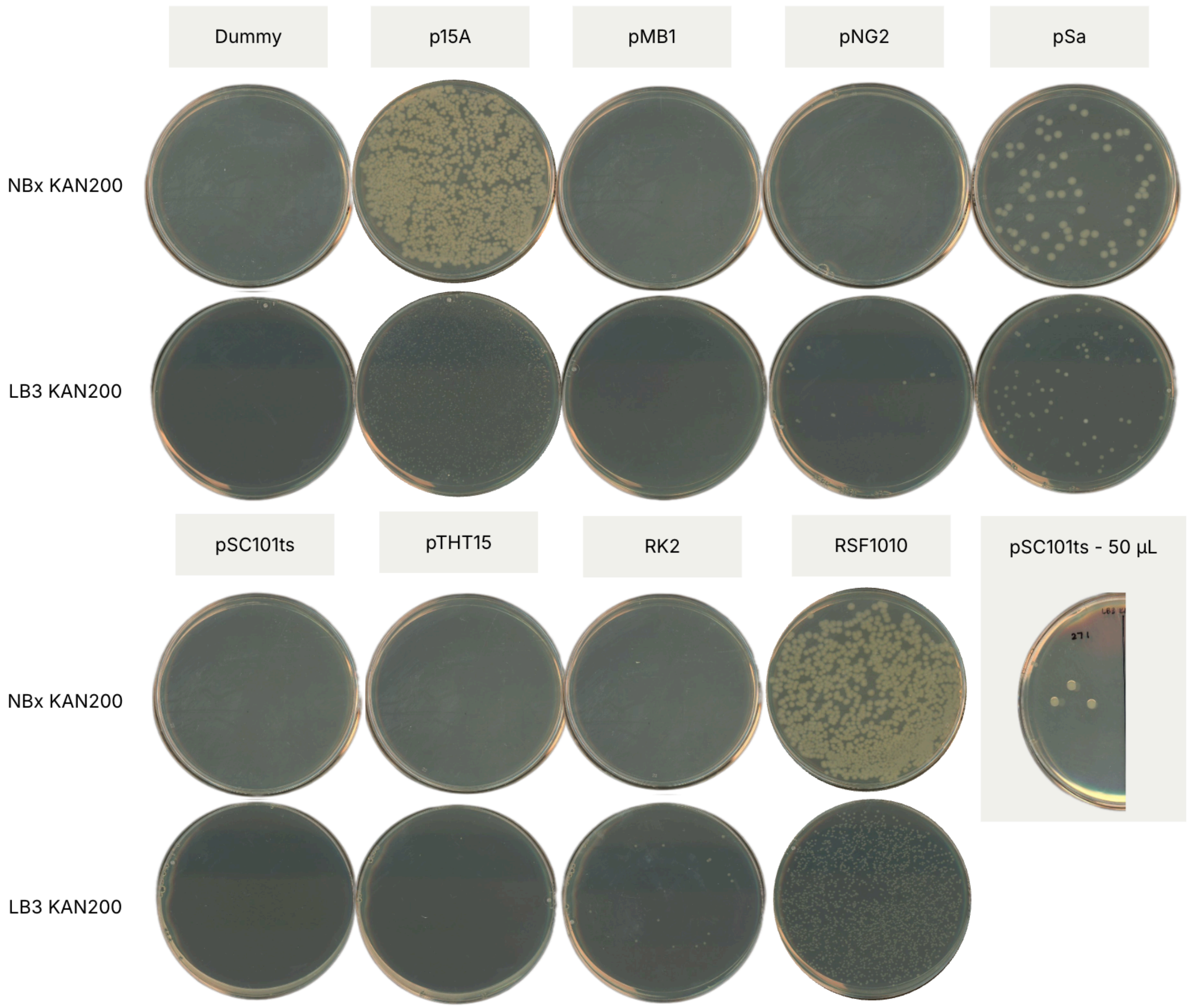


**Supplementary Figure 4. Agar selection of individual plasmids for *V. natriegens* NBx CyClone™.** Images of selective agar plates for *V. natriegens* NBx CyClone™ following transformation with individual plasmids (see Supplementary Table 3). Transformants were selected on both NBx media and LB3 media. Kanamycin concentrations are reported in µg/mL. Note: transformants for pSC101ts plasmid were only visible after plating 50 µL of the transformation on LB3 KAN200.

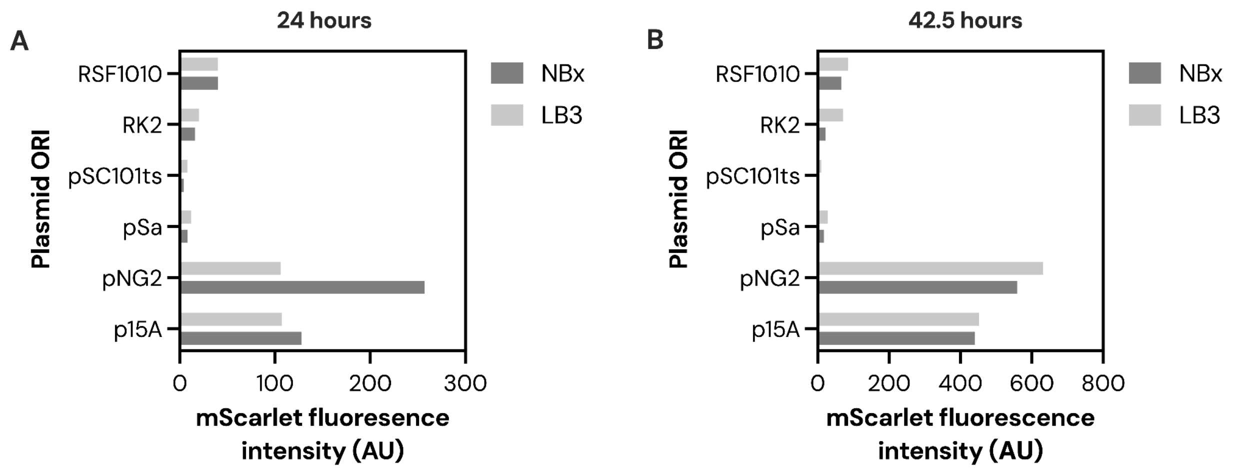


**Supplementary Figure 5. mScarlet fluorescence from *V. natriegens* NBx CyClone™.** mScarlet fluorescence intensity from individual plasmids over **(A)** 24 and **(B)** 42.5 hours in NBx and LB3 media.

### Supplementary Tables

**Supplementary Table 1. Plasmids and plasmid libraries used in this study.** Parts for building these plasmids by Golden Gate assembly can be found through Addgene Kit #1000000234. The 23 ORIs used in the plasmid libraries can be seen in Figure 1.

| Plasmid or library name | Origin of replication | Selective marker | Source |
|---|---|---|---|
| Plasmid Libraries | | | |
| pGL2_217 | 23 ORIs | kanamycin | [1] |
| pGL2_218 | 23 ORIs | gentamicin | [1] |
| pGL2_219 | 23 ORIs | erythromycin | [1] |
| pGL2_220 | 23 ORIs | tetracycline | [1] |
| pGL2_221 | 23 ORIs | chloramphenicol | [1] |
| pGL2_222 | 23 ORIs | spectinomycin | [1] |
| Individual Plasmids | | | |
| pGL2_228 | Dummy | kanamycin | [1] |
| pGL2_250 | p15A | kanamycin | [1] |
| pGL2_268 | pMB1 | kanamycin | [1] |
| pGL2_269 | pNG2 | kanamycin | [1] |
| pGL2_270 | pSa | kanamycin | [1] |
| pGL2_271 | pSC101ts | kanamycin | [1] |
| pGL2_273 | pTHT15 | kanamycin | [1] |
| pGL2_276 | RK2 | kanamycin | [1] |
| pGL2_277 | RSF1010 | kanamycin | [1] |
| pUC19 | pMB1 derivative | carbenicillin | [2] |

**Supplementary Table 2. Antibiotic concentrations tested in this study.** Antibiotic concentrations tested for liquid and agar selection of NBx CyClone™ strain using the POSSUM assay. Bolded values indicate concentrations where transformants were observed and used for downstream NGS analysis.

| Antibiotic | Liquid selective concentration (µg/mL) | | | | Agar selective concentration (µg/mL) | Plasmid or plasmid library |
|---|---|---|---|---|---|---|
| | 0.25X | 1X | 4X | 6X | | |
| KAN | 12.5 | 50 | 200 | **300** | **200** | pGL2_217 |
| GEN | 5 | 20 | 80 | | **80** | pGL2_218 |
| ERM | 1 | 4 | 16 | | 16 | pGL2_219 |
| TET | 2.5 | **10** | 40 | | **40** | pGL2_220 |
| CAM | **8.5** | 34 | 136 | | **5** | pGL2_221 |
| SPEC | 12.5 | 50 | 200 | | 12.5 | pGL2_222 |
| CARB | 12.5 | **50** | 200 | | **50** | pUC19 |

**Supplementary Table 3. Transformation efficiency of individual plasmids in *V. natriegens* NBx CyClone™.**

| Plasmid | ORI | Functional (Y/N) | Transformation efficiency (CFU/µg DNA) |
|---|---|---|---|
| pGL2_228 | Dummy | N | 0 |
| pGL2_250 | p15A | Y | $5.7 \times 10^{7}$ |
| pGL2_268 | pMB1_*B. longum* | N | 0 |
| pGL2_269 | pNG2 | Y | $2.5 \times 10^{5}$ |
| pGL2_270 | pSa | Y | $1.4 \times 10^{7}$ |
| pGL2_271 | pSC101ts | Y | $1.3 \times 10^{3}$ |
| pGL2_273 | pTHT15 | N | 0 |
| pGL2_276 | RK2 | Y | $3.1 \times 10^{5}$ |
| pGL2_277 | RSF1010 | Y | $6.2 \times 10^{7}$ |

**Supplementary Table 4. Primers used in this study.** Regions of homology to the fragment being amplified are shown in bold. The underlined regions in PCR 1 primers represent 5' primer extensions which introduce stub sequences enabling binding of dual indexed Illumina i5 and i7 primers in the subsequent PCR 2 step. A mix of primers containing zero to six random bases (N) was used to add diversity to the amplicon library for sequencing on a MiniSeq.

| Primer | Sequence | Description |
|---|---|---|
| oCG0038 | TCCCTACACGACGCTCTTCCGATCT[NNNNNN]**GGGTCACGCGTAGGACG** | PCR 1 forward primer for amplification of the plasmid barcode |
| oCG0039 | GAGTTCAGACGTGTGCTCTTCCGATCT[NNNNNN]**CCAGCTTCACACGGCGT** | PCR 1 reverse primer for amplification of the plasmid barcode |

### References


1. Gilbert, C. *et al.* Design and construction towards a pan-microbial toolkit. *bioRxiv* 2024.02.23.581749 (2024) doi:10.1101/2024.02.23.581749.
2. *NBx CyClone™ Product Manual*. (2025).
